# Improving Turnaround Times with Artificial Intelligence in Microbiology

**DOI:** 10.64898/2026.03.09.710721

**Authors:** Ross Davidson, Charles Heinstein, Glenn Patriquin, Lee William Goneau, Leah Ashleigh Brown, Brian James Hill

## Abstract

This dual-center study evaluated the impact of artificial intelligence (AI) on urine culture turnaround times in Canadian diagnostic laboratories employing full microbiology laboratory automation. Data were collected before and after the implementation of PhenoMATRIX (PM), an AI-based software designed to support culture sorting and result interpretation. In both a low-volume tertiary care hospital and a high-volume community laboratory, PM reduced the time to final culture reporting, with decreases of approximately 1.5 hours and 3.9 hours, respectively. Implementation of PM+, which automatically releases defined results to patient charts, further improved turnaround time. These findings indicate that microbiology laboratories with full laboratory automation can achieve further improvements in turnaround time by integrating AI-culture assessment and results release.

## INTRODUCTION

Automation in clinical microbiology has become essential in response to persistent staffing shortages and increasing specimen volumes. Automated technologies improve efficiency compared to manual methods across laboratories of varying size and patient acuity.^1^ Following implementation of automation, artificial intelligence (AI) algorithms offer the next step in optimizing workflows.

Copan’s PhenoMATRIX (PM) software (Copan Diagnostics, Murrieta, CA, 92652) uses AI algorithms to assess and sort culture plates based on software interpretations of bacterial growth from digital plate images. For urine cultures, the software categorizes plates demonstrating no growth, no significant growth (< 10 colonies/plate), mixed growth (> 2 colony types), and/or specific colony colorations on chromogenic media. Prior studies have shown the accuracy of these algorithms across multiple specimen types and media.^2-6^ Beyond accuracy, AI has the potential to accelerate result reporting through continuous plate assessment.

This study evaluated time to result reporting (TTRR) using PM in a dual-center Canadian study of urine cultures. In addition, PhenoMATRIX PLUS (PM+), a feature that automatically releases defined culture results to the patient chart without staff intervention, was evaluated at one site. This represents the first dual-site evaluation of PM AI-driven algorithms in North America.

## METHODS

This study was conducted at two Canadian diagnostic laboratories with differing volumes, patient populations, and geographic settings. The first site, Queen Elizabeth II Health Sciences Centre (QEII), is a large tertiary care hospital located in Halifax and provides diagnostic testing on behalf of the Nova Scotia Health Authority for other laboratories across the province. The laboratory also serves as the reference laboratory for the Canadian Atlantic region, processing over 65,000 urine specimens annually with volumes increasing by approximately 10% each year. The second site, Dynacare Laboratories in Brampton, Ontario, is a high-volume community laboratory operating 24/7 and processing an annual average of approximately 650,000 urine specimens collected across the province.

TTRR data were collected before PM implementation (PrePM) and after full software implementation (PostPM; including PM+ at Halifax). For Dynacare, PrePM and PostPM TTRR were defined as the time from specimen placement on the WASPLab (Copan Diagnostics, Murrieta, CA, 92652) to final approval and release of the culture report in the laboratory information system (LIS). At QEII, TTRR was calculated from specimen receipt in the laboratory to result release in the LIS.

Urine specimens were plated and incubated using WASPLab automation and imaged after 16 hours of incubation. The study was conducted over a 3-year period from May 2022 to May 2025. Over the study period, no other significant changes (e.g., staffing, workflow, process) aside from PM implementation were made in either lab that the authors believe could confound the results presented.

At QEII, urine cultures were screened on the day shift (07:00 – 16:00) for all TTRR timepoints measured (PrePM, PostPM, and PM+). Urine cultures with a final result of no growth or normal urogenital flora of less than 10 colonies were automatically released and reported using PM+. At Dynacare, urine culture screening was performed on the evening shift (16:00 – 24:00) during PrePM data collection, and during the day and evening shifts (08:00 – 24:00) for PostPM data collection. PrePM, PostPM, and PM+ data collection periods, and media used for urine culture testing is listed in Table 1:

**Table 1:**
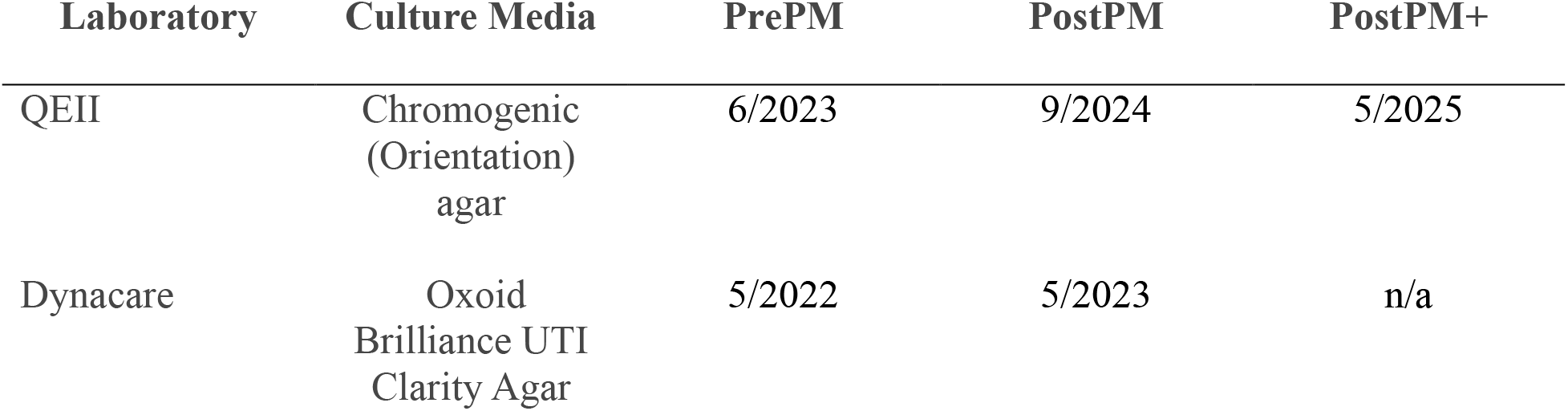
Culture medium and Date of PhenoMatrix implementations.

| Laboratory | Culture Media | PrePM | PostPM | PostPM+ |
| --- | --- | --- | --- | --- |
| QEII | Chromogenic<br>(Orientation)<br>agar | 6/2023 | 9/2024 | 5/2025 |
| Dynacare | Oxoid<br>Brilliance UTI<br>Clarity Agar | 5/2022 | 5/2023 | n/a |

## RESULTS

### QEII

During the study period the daily average of urine cultures increased from 167 in 2023 to approximately 180 in 2024-2025, representing an ∼7% volume increase. The TTRR for PrePM, PostPM and PostPM+ phases are shown in Figure 1.

**Figure 1.**
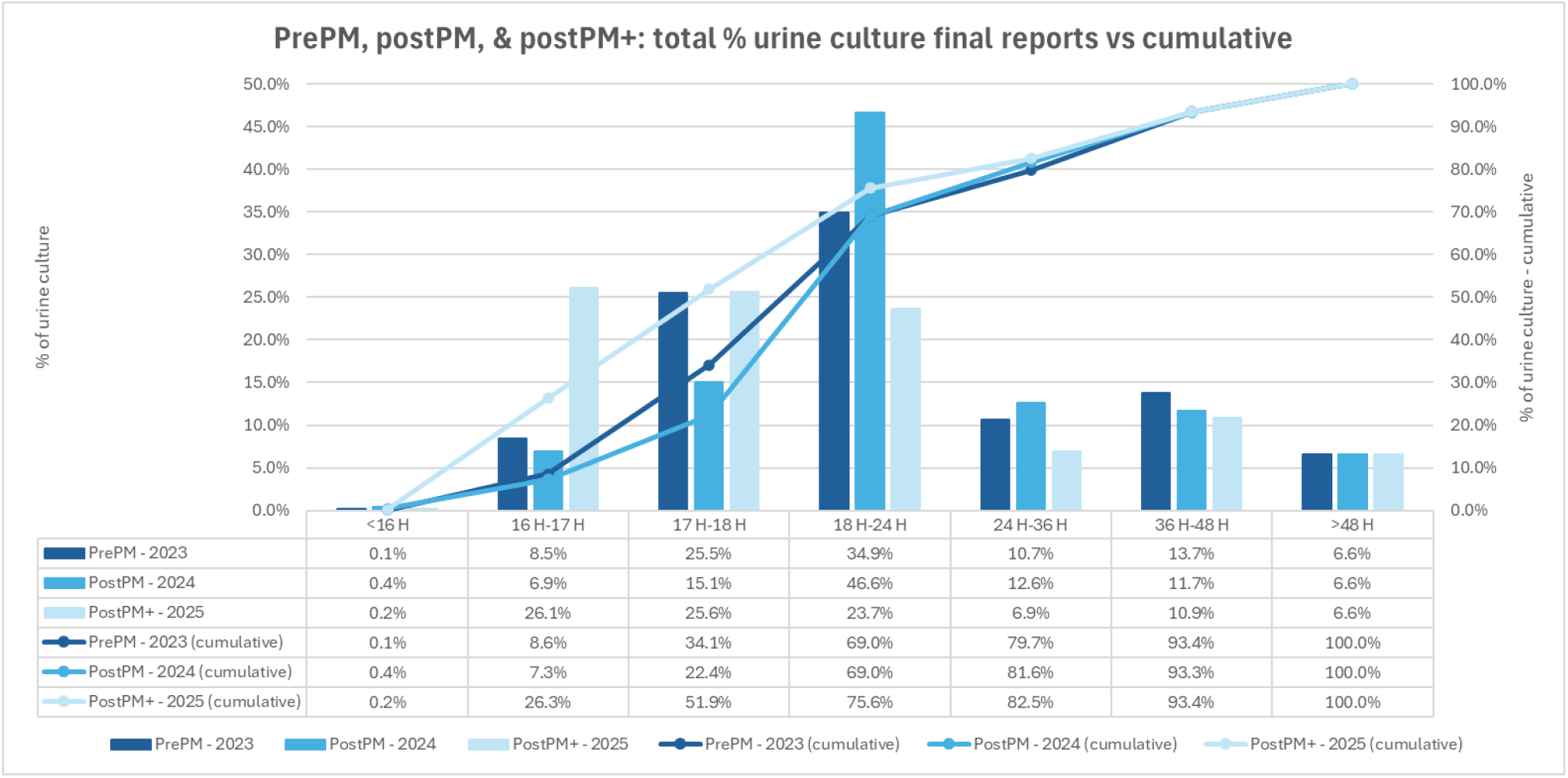
TTRR for PrePM, PostPM and PostPM+ final culture reports, QEII.

With the implementation of PM+, the most significant increase in final reports released occurred within the first two hours after the 16-hour image was taken. The percentage of final reports released by 17 hours increased to 26.3%, compared with 8.6% in the PrePM phase and 7.3% in PostPM. By 18 hours, 51.9% of urine cultures had a final report available, compared with 34.1% (PrePM) and 22.4% (PostPM). At 24 hours, 75.6% of final urine culture results were released with PM+, compared with 69.0% in both PrePM and PostPM phases. Of note, approximately 77% of the QE II’s urine cultures are negative or reported as normal urogenital flora.

Consequently, the proportion of cultures pending final reports beyond 24 hours decreased from 31.0% (PrePM and PostPM) to 24.4% (PM+). Overall, the mean TTRR decreased by approximately 1 hour and 26 minutes between June 2023 and May 2025.

### Dynacare

At Dynacare, the average daily urine culture volume increased from 1593 in 2022 to 1778 in 2023 (∼ 11.6% volume increase). The percentage of negative urine cultures reported as no growth (NG) or non-significant growth (NSG) accounted for 53.18% of overall urine cultures in May 2022 (prePM) and 50.88% in May 2024 (postPM). TTRR for PrePM and PostPM phases are shown in Figure 2.

**Figure 2.**
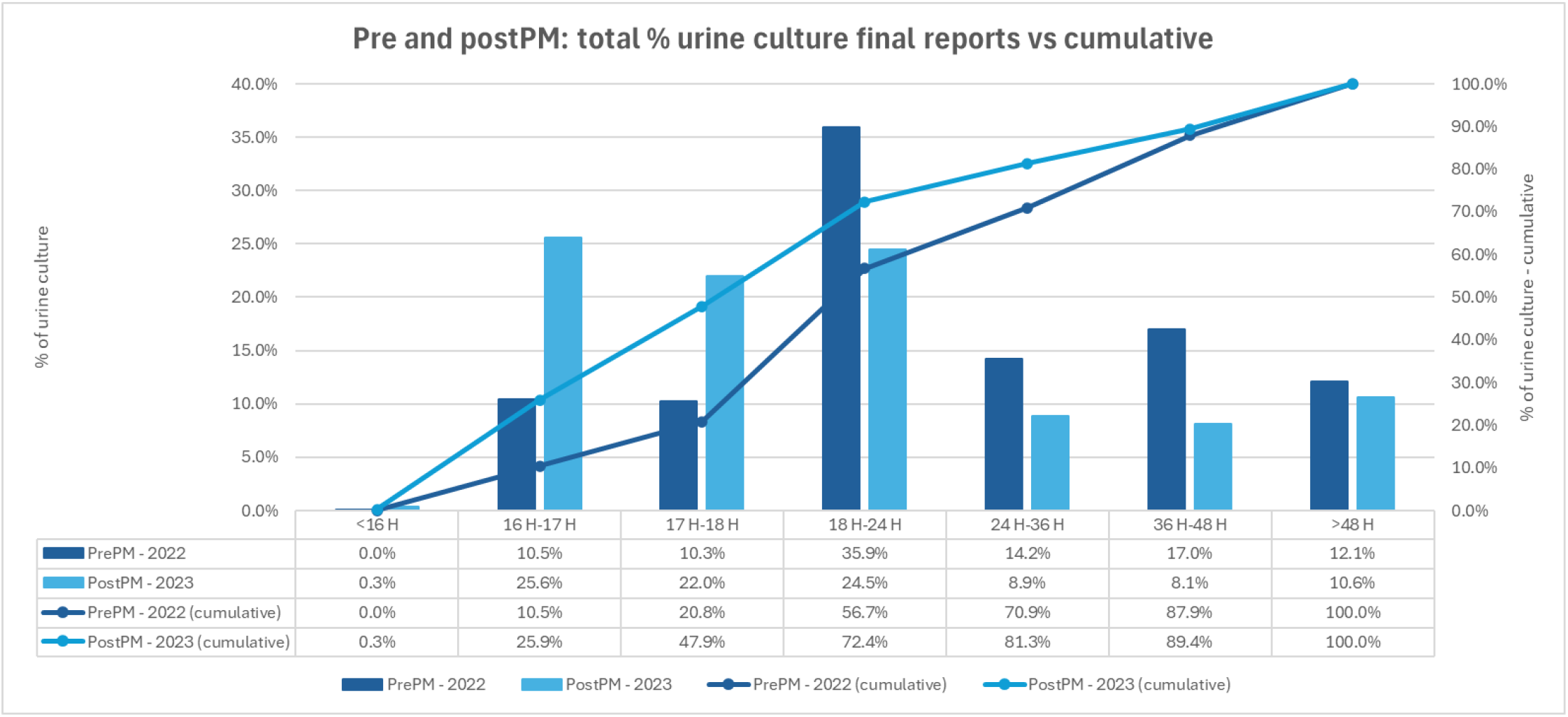
TTRR results for PrePM and PostPM final culture reports; Dynacare.

Following PM implementation, a higher proportion of urine culture final reports were released within 18 hours compared to those released after 18 hours. The proportion of final culture reports released by 17 hours increased from 10.5% (PrePM) to 25.9% (PostPM). By 18 hours, 47.9% of reports were finalized compared to 20.8% before implementation, and by 24 hours, 72.4% of final urine culture reports were released versus 56.7% PrePM. The proportion of final culture reports pending beyond 24 hours decreased by 15.7% (43.3% PrePM to 27.6% PostPM). Overall, mean final report TTRR improved by approximately 3 hours and 51 minutes from May 2022 to May 2023.

## DISCUSSION

This dual-site study demonstrates that artificial intelligence can significantly enhance efficiency in microbiology laboratories using full automation, regardless of laboratory size or specimen volume. Both the tertiary care hospital and community laboratory settings achieved meaningful reductions in TTRR following the implementation of PhenoMATRIX algorithms. The QEII laboratory achieved an average TTRR reduction of approximately 1.5 hours, while Dynacare observed nearly a 4-hour reduction.

These findings align with previous work by Culbreath et al., which reported accelerated reporting times for positive cultures and a decreasing volume of cultures reported at longer time periods when algorithm-assisted reporting was used. The current data extends these observations by confirming that AI-driven plate interpretation can improve turnaround times in both hospital-based and high-volume community laboratories, even when baseline workflows and throughput differ substantially.

The ongoing shortage of qualified medical laboratory technologists and increasing specimen volumes underscores the importance of technologies that optimize efficiency without compromising quality. Full laboratory automation has already been shown to improve culture processing and mitigate staffing challenges. ^7-9^ The addition of AI software such as PhenoMATRIX builds on these advantages by enabling continuous digital analysis of culture plates and earlier result release.

At QEII, the introduction of PM+, which automatically releases defined negative or normal flora results directly to patient charts, further enhanced workflow efficiency. By reducing the need for manual review of non-significant cultures, laboratories can redirect technologist expertise to more complex tasks, thereby improving overall productivity and patient care.

The combined benefits of continuous incubation, automated imaging, and AI-assisted interpretation represent a major advancement in clinical microbiology. Smart incubation has been shown to shorten incubation times while maintaining accuracy and enabling faster identification and antimicrobial susceptibility testing results.^10-12^ Future innovations, such as AI-assisted selection of pathogens for automated identification and susceptibility testing could enable end-to-end automation of uncomplicated cultures. This evolution will allow laboratory professionals to focus their expertise on clinically complex specimens while maintaining high throughput and timely patient results.

This study is, to our knowledge, the first dual-site evaluation in North America assessing the effects of PM AI-driven algorithms on urine culture turnaround times. These findings demonstrate that AI integration into automated microbiology workflows can produce measurable improvements in TTRR across diverse laboratory environments. The incorporation of auto-release features such as PM+ further enhances these gains, offering a promising pathway toward fully automated, efficient, and clinically responsive diagnostic microbiology.

